# Comparative analysis of muscle thickness, muscle percentage of subcutaneous tissue, and muscle luminance in skeletal muscles of limbs of normal children and children with limb disabilities due to cerebral palsy using ultrasound imaging

**DOI:** 10.1101/2024.06.24.600485

**Authors:** Hideki Ishikura

## Abstract

**Background:** Ultrasound imaging is increasingly utilized for musculoskeletal evaluation because it is cost-effective and non-invasive. This study aims to analyze the differences in muscle thickness, muscle percentage of subcutaneous tissue, and muscle luminance in children with limb disabilities due to cerebral palsy compared with typically developing children.

**Methods:** This cross-sectional study included 12 children aged 6–15 years, divided equally into two groups: children with physical disabilities due to cerebral palsy and healthy controls. Ultrasound imaging was used to measure muscle thickness, percentage of muscle in subcutaneous tissue, and muscle luminance in various muscle groups of the limbs. Statistical analyses were conducted using t-tests to compare the two groups.

**Results:** Significant differences were observed in muscle thickness and percentage of muscle in subcutaneous tissue between the groups, particularly in the lower limbs. Muscle luminance did not exhibit consistent differences. These findings suggest a decreased muscle mass and altered composition in children with cerebral palsy, which may influence their physical function and mobility.

**Conclusions:** Our results underscore the value of ultrasound in clinically assessing muscle properties in children with cerebral palsy. This modality offers a promising tool for evaluating muscle alterations and potentially guiding targeted interventions to improve mobility and quality of life in affected children.

## Introduction

In rehabilitation, evaluating the musculoskeletal system is crucial to establishing an appropriate treatment plan to restore patient function. This evaluation requires a detailed understanding of the structures of muscles, tendons, joints, and ligaments, as well as the degree of their function and impairment. In the older adults, muscle quantity and quality are important factors in maintaining physical function [1]. Muscle quality is assessed by magnetic resonance imaging (MRI) and ultrasound imaging, including the amount and distribution of fat within the muscle and the shear rate of the muscle [2]. In addition to tests and measurements performed by physical therapists, including manual muscle strength testing and joint range of motion measurements, musculoskeletal evaluation also includes imaging tests such as X-rays, computed tomography (CT) scans, and MRI. Imaging tests provide a detailed picture of the musculoskeletal structures and are effective in determining the degree of function and disability. However, these imaging techniques have limitations, such as being expensive and requiring the use of radiation to the patient. Nevertheless, evaluation of the musculoskeletal system using ultrasound imaging has gained attention in recent years. Ultrasound imaging offers real-time visualization of structures within the body, is less invasive to the patient, and can be performed at a relatively low cost. Furthermore, an increasing number of ultrasound diagnostic imaging systems are portable and can be used in various environments, such as at the bedside or in home settings. With this advancement, various studies have been conducted using ultrasound imaging to evaluate the musculoskeletal system. In a previous study that investigated muscle mass in the elderly, the thickness of the rectus femoris muscle was measured using ultrasound images and compared with body composition and mobility function; muscle thickness assessment using ultrasound images was associated with muscle mass [3]. Muscle luminance, which is derived from the echo intensity of images in ultrasound imaging, is negatively correlated with physical functions such as standing movements and walking speed [4]. Other previous studies examining the relationship between physical function and ultrasound images in patients with chronic kidney disease indicated that intramuscular fat and fibrotic tissue are factors leading to poor physical function [5]. These studies demonstrate that ultrasound imaging can be used to evaluate skeletal muscle quantitatively and qualitatively, assessing aspects such as connective tissue within the muscle and its relationship with physical function and muscle strength, making it a valid evaluation in clinical practice.

In pediatric rehabilitation, motor function assessments such as Gross Motor Function Measure and Gross Motor Function Classification System (GMFCS), activities of daily living such as Wee Functional Independence Measure, and other assessments of activities of daily living are commonly used. Musculoskeletal evaluation using ultrasound imaging in children has also been conducted, with previous studies reporting that muscle thickness correlates with sprinting ability and physical activity. Secondary sarcopenia can occur in children as well as in adults because of reduced skeletal muscle mass, especially with decreased activity levels [6]. Reports of ultrasound evaluation of skeletal muscles of the limbs in children are available; however, few reports examine children with limb dysfunction due to cerebral palsy or other causes. This is because in evaluating children with severe physical disabilities, many evaluation tasks cannot be performed due to differences in intellectual function and development. Consequently, few quantitative evaluations can be performed on both mildly and severely disabled patients. Evaluating children with physical disabilities using ultrasound imaging is effective in clinical practice because it allows for the assessment of other dynamic aspects of the child and their relationship with the child’s ability to perform activities of daily living.

In this study, we report our findings by evaluating skeletal muscles using ultrasound imaging and examining the relationship between movement performance and skeletal muscle structure in healthy individuals and children/people with physical disabilities due to cerebral palsy.

## Materials and Methods

The participants included six healthy children and six children with physical disabilities due to cerebral palsy, aged 6–15 years. The ages of the healthy children and those with disabilities were 8.7 ± 9.8 years and 10.8 ± 2.3 years, respectively. The recruitment period for the study was from May 23, 2023, to March 31, 2024.

The children with physical disabilities were classified by GMFCS based on their level of mobility function and intellectual development as “E: able to perform simple calculations / D: able to understand simple letters and numbers / C: able to understand simple colors and numbers / B: able to understand simple language / A: unable to understand language.”

An ultrasound imaging device (FAMUBO-W, Seiko) was used for subcutaneous morphometric measurements. During the measurement, participants were instructed to relax in the supine position, and images in the short-axis direction were taken at the maximum bulge of the skeletal muscle being evaluated. Images were taken on the left and right sides, and the average value was used as the result. For the measurement, a sufficient amount of ultrasonic coupling agent was applied to the skin surface, and the ultrasonic probe was brought into contact with the skin without applying pressure, following the method of a previous study [7]. The images were analyzed for muscle thickness and muscle luminance using image analysis software (Image J, National Institutes of Health). Muscle thickness was evaluated by measuring the distance from the bone index to the fascia at a position perpendicular to the bone index for the elbow flexor muscle group (anterior surface of the upper arm), wrist flexor muscle group (anterior surface of the forearm), knee extensor muscle group (anterior surface of the thigh), and ankle plantar flexor muscle group (posterior surface of the lower leg) (Figure 1).

**Fig. 1.**
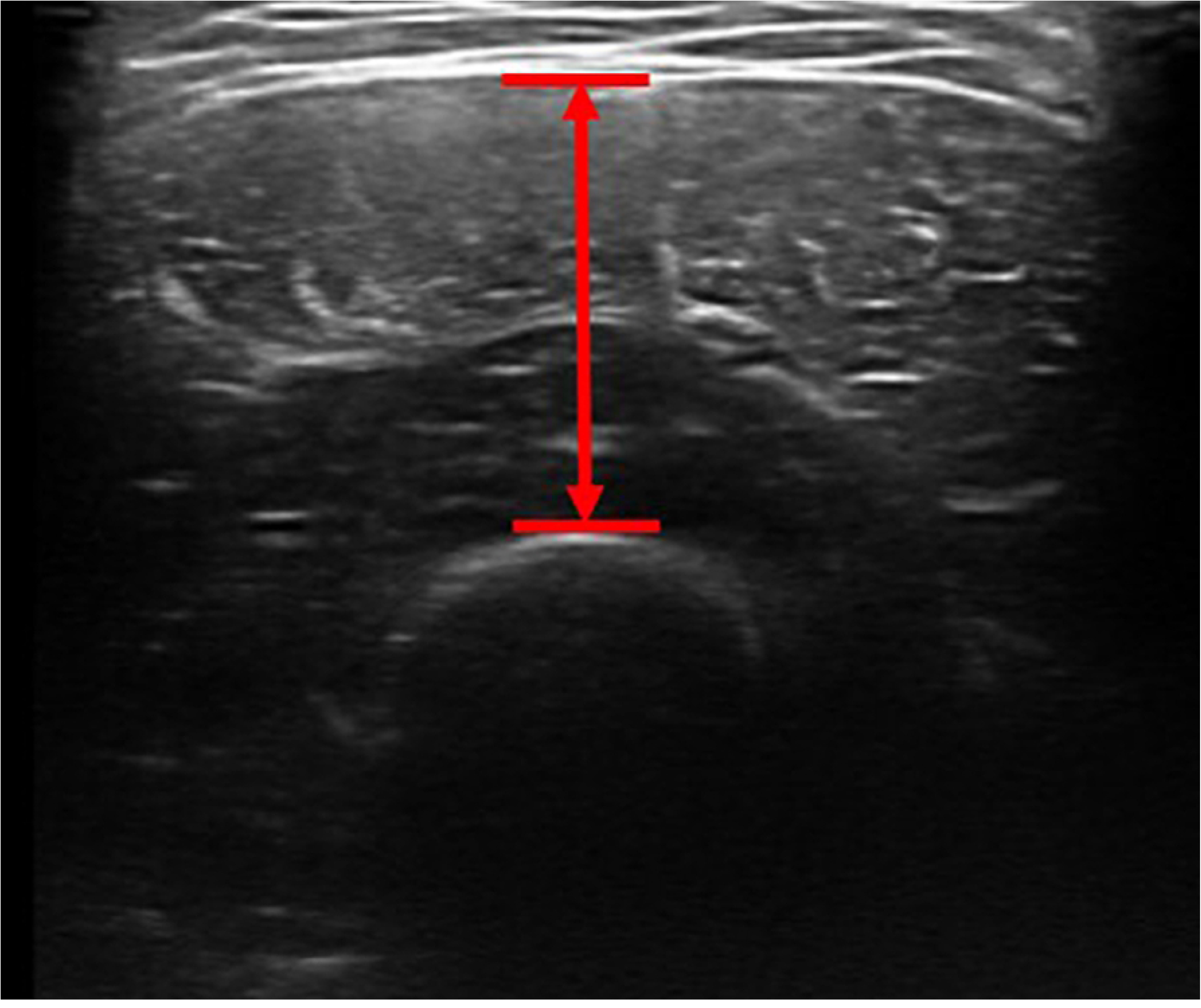
Measurement of muscle thickness Measure the distance from the bone index to the fascia at a position perpendicular to the bone index

The distance from the bone index to the skin surface was also measured to determine the percentage of muscle in the entire subcutaneous tissue. For muscle luminance, we specified the largest possible rectangular area for skeletal muscle, avoiding fascia, and used an 8-bit grayscale with 256 shades (0–255) within the specified area (Figure 2). In this evaluation, higher values indicate higher muscle luminance, suggesting that connective tissues other than myofibers are contained within the myoplasm [7].

**Fig. 2.**
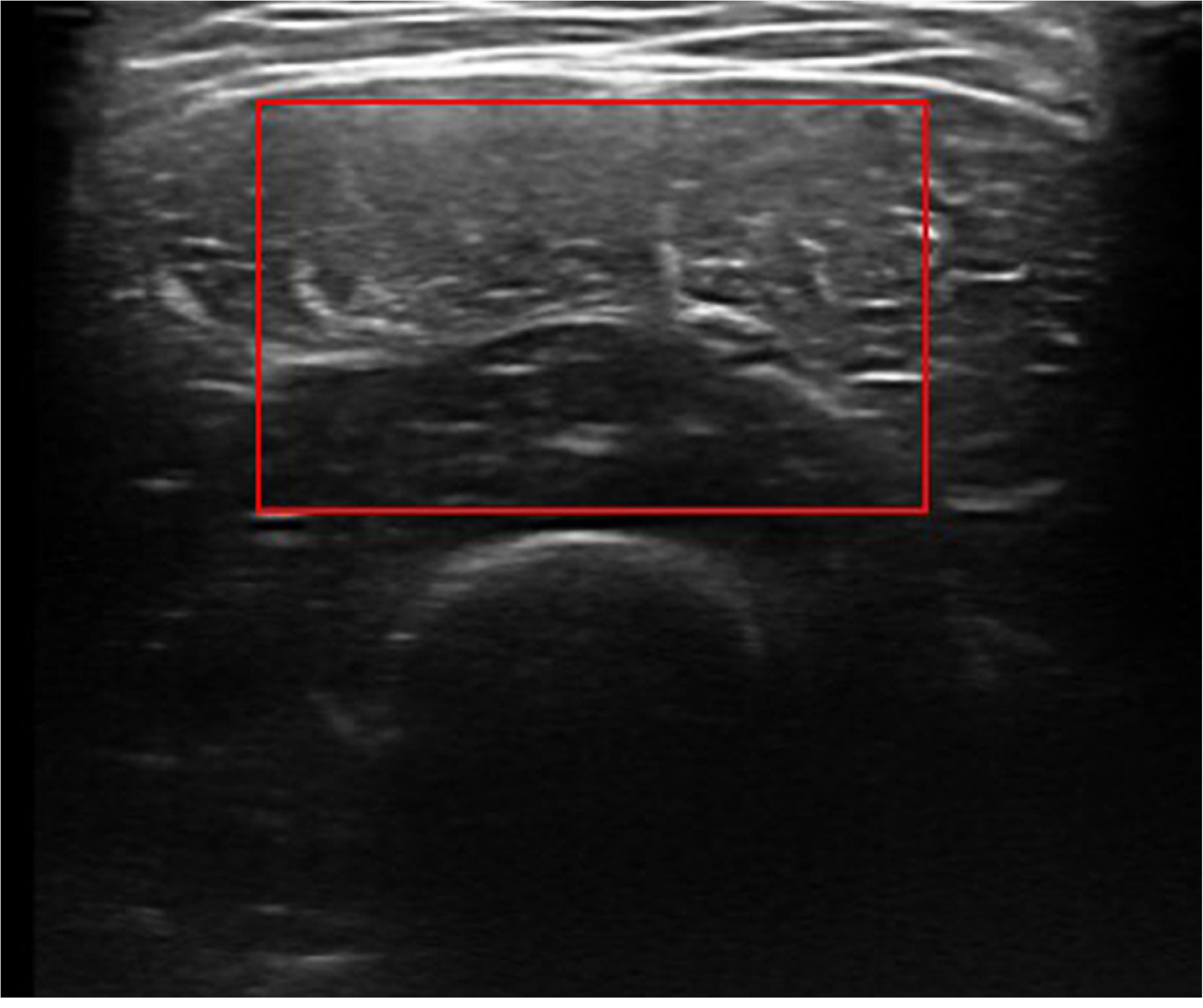
Measurement of muscle luminance

Designate the largest possible rectangular area for skeletal muscle, avoiding fascia. Statistical analysis was performed using statistical software (SPSS Statistics, IBM). The data was evaluated for normality with the Shapiro–Wilk test and confirmed to follow a normal distribution (p > 0.05). Comparisons between healthy children and children with physical disabilities due to cerebral palsy were made using t-tests for muscle thickness, percentage of muscle in the total subcutaneous tissue, and muscle luminance of the elbow flexor muscle group (front of the upper arm), wrist flexor muscle group (front of the forearm), knee extensor muscle group (front of the thigh), and ankle plantar flexor muscle group (back of the lower leg). The significance level was set at 5%.

The Research Ethics Committee of Hiroshima Cosmopolitan University (Approval No. 2023003) approved this study. Following the Declaration of Helsinki, this study was conducted in consideration of ethics and personal information, and the research content was explained in writing and orally to the affiliated institution, the participants, the participant’s guardian, and key persons as surrogates. Written informed consent was obtained before the study was conducted.

## Results

The mobility of the participants with physical disability in this study was more limited than that of the healthy ones, with GMFCS of II:1, III:1, IV:1, and V:3. Intellectual development was D: 1, B: 2, and A: 3, with half of them unable to understand language. The results (mean ± standard deviation) for the elbow flexor muscle group on the anterior surface of the upper arm are presented in Table 1. The muscle thickness of the elbow flexor group was 19.16 ± 1.99 mm in the healthy children and 16.08 ± 6.11 mm in the children with physical disabilities due to cerebral palsy, with no significant difference between the groups (p=0.285). The percentage of muscle in the total subcutaneous tissue of the anterior surface of the upper arm was 88.20 ± 3.11 % in the healthy children and 77.62 ± 7.58 % in the children with physical disabilities due to cerebral palsy, with a significant difference between the groups (p=0.01). Muscle luminance of the elbow flexor group was 83.85 ± 6.07 % in the healthy children and 80.69 ± 28.48 % in the children with physical disabilities due to cerebral palsy, with no significant difference between the groups (p=0.796). These results indicate that the elbow flexor group had a decreased percentage of muscle in the overall subcutaneous tissue of the upper arm. Additionally, a trend toward decreased muscle mass in the limb flexors was observed, although with no significant difference in muscle thickness.

**Table 1:**
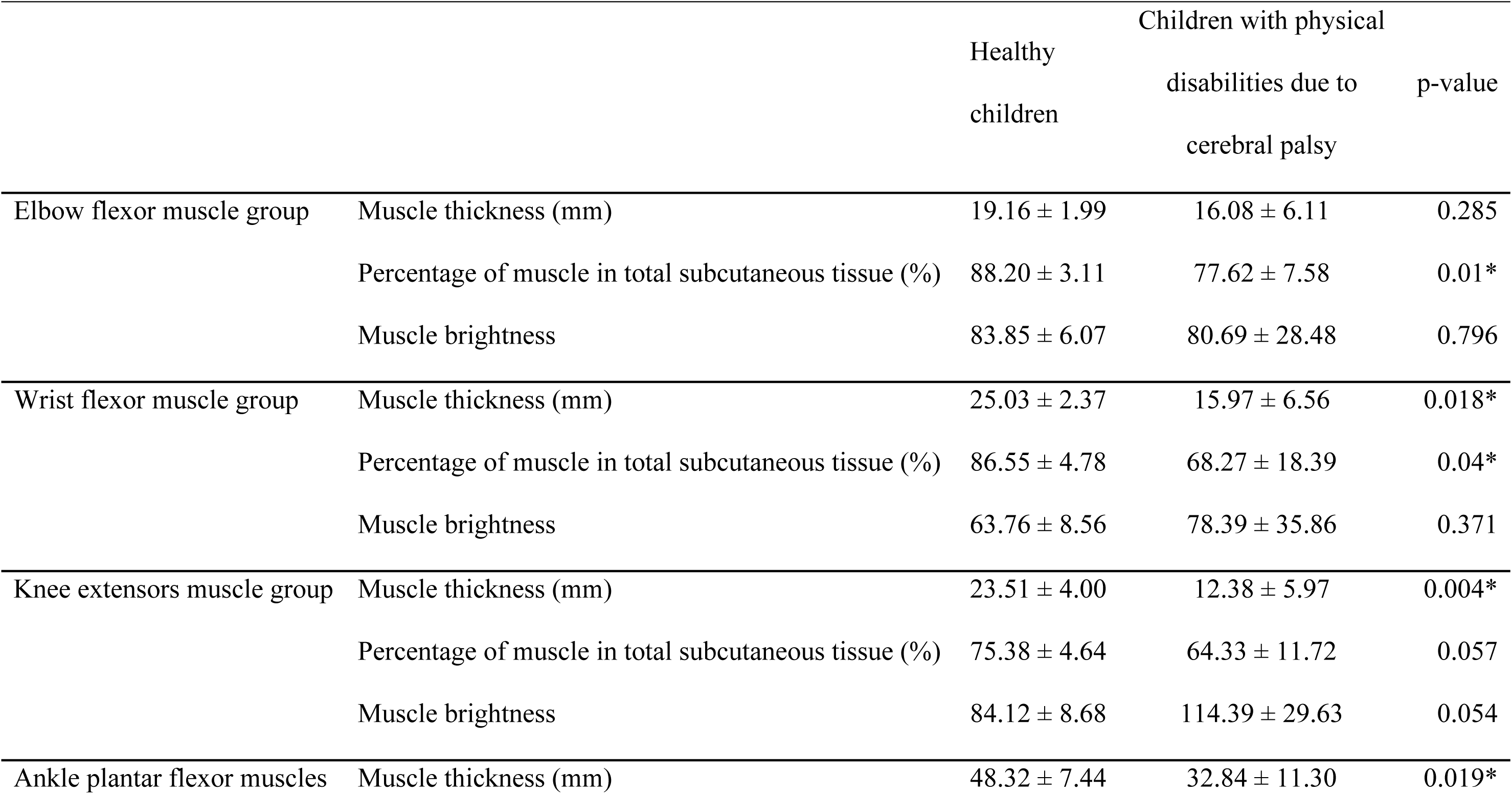
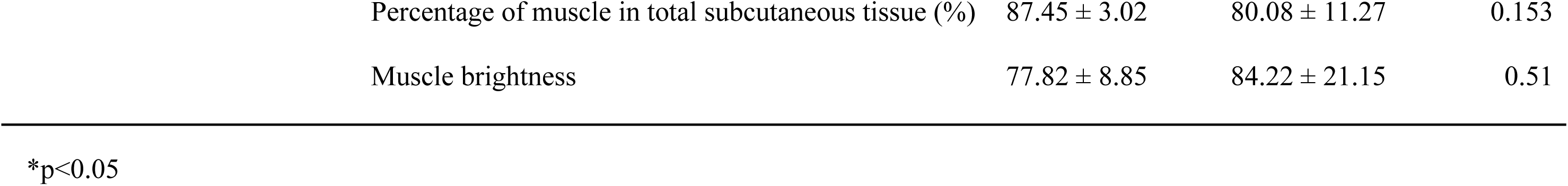
Results of analysis of body morphology using ultrasound images.

The results (mean ± standard deviation) for the wrist flexor muscle group on the anterior surface of the forearm are presented in Table 1. The muscle thickness of the wrist flexor group was 25.03 ± 2.37 mm in the healthy children and 15.97 ± 6.56 mm in the children with physical disabilities due to cerebral palsy, with a significant difference between the groups (p=0.018). The percentage of muscle in the total subcutaneous tissue of the anterior forearm was 86.55 ± 4.78 % in the healthy children and 68.27 ± 18.39 % in the children with physical disabilities due to cerebral palsy, with a significant difference between the groups (p=0.04). The muscle luminance of the wrist flexor group was 63.76 ± 8.56 % in the healthy children and 78.39 ± 35.86 % in the children with physical disability, with no significant difference between the groups (p=0.371). These results indicate that in the forearms, children with physical disabilities due to cerebral palsy had decreased muscle mass and percentage of muscle in the total subcutaneous tissue than healthy children.

The results (mean ± standard deviation) for the knee joint extensor muscle group on the anterior aspect of the thigh are presented in Table 1. The muscle thickness of the knee extensor muscle group was 23.51 ± 4.00 mm in the healthy children and 12.38 ± 5.97 mm in the children with physical disabilities due to cerebral palsy, with a significant difference between the groups (p=0.004). The percentage of muscle in the total subcutaneous tissue of the anterior thigh was 75.38 ± 4.64 % in the healthy children and 64.33 ± 11.72 % in the children with physical disabilities due to cerebral palsy, with no significant difference between the groups (p=0.057). Muscle luminance of the knee extensor group was 84.12 ± 8.68 % in the healthy children and 114.39 ± 29.63 % in the children with physical disabilities due to cerebral palsy, with no significant difference between the groups (p=0.054). These results indicated that in the thighs, muscle thickness was reduced in the children with physical disabilities due to cerebral palsy. No significant difference was observed in the percentage of muscle in the overall thigh subcutaneous tissue of the children with physical disabilities due to cerebral palsy. The p-value was close to the significance level, indicating a tendency for the percentage of muscle to decrease in these children. Similarly, no significant difference was observed in the muscle luminance of the knee extensor muscle group, but the p-value was close to the significance level, and muscle luminance tended to increase in children with physical disabilities due to cerebral palsy.

The results (mean ± standard deviation) for the ankle plantar flexor muscle group on the posterior aspect of the lower leg are presented in Table 1. The muscle thickness of the ankle plantar flexor group was 48.32 ± 7.44 mm in healthy children and 32.84 ± 11.30 mm in children with physical disabilities due to cerebral palsy, with a significant difference between the groups (p=0.019). The percentage of muscle in the total subcutaneous tissue of the posterior aspect of the lower leg was 87.45 ± 3.02 % in the healthy children and 80.08 ± 11.27 % in the children with physical disabilities due to cerebral palsy, with no significant difference between the groups (p=0.153). Muscle luminance of the ankle plantar flexor group was 77.82 ± 8.85 in the healthy children and 84.22 ± 21.15 in the children with physical disabilities due to cerebral palsy, with no significant difference between the groups (p = 0.51). These results revealed that in the lower leg, muscle thickness decreased in children with physical disabilities due to cerebral palsy.

## Discussion

This study investigated the relationship between movement performance and skeletal muscle structure by evaluating the musculoskeletal system using ultrasound imaging in healthy children and children with physical disabilities due to cerebral palsy aged 6 to 15 years. The severity classification of the participants with physical disabilities was II:1, III:1, IV:1, and V:3 in GMFCS, indicating they had limited mobility function compared with the healthy participants. The results of this study revealed a decrease in the percentage of muscle in the total subcutaneous tissue in the upper extremities and a marked reduction in muscle thickness in the lower extremities.

Regarding the muscles of the upper limb, no significant differences were observed between the non-disabled children and children with physical disabilities due to cerebral palsy in the muscle thickness of the elbow flexor group. However, non-disabled children exhibited a significantly higher percentage in the muscle thickness of the forearm flexor group and total subcutaneous tissue of the elbow flexor group and forearm flexor group than the children with physical disabilities due to cerebral palsy. This indicates that children with physical disabilities due to cerebral palsy have decreased muscle mass, especially relative muscle mass in the subcutaneous tissue, which may be related to decreased frequency of upper extremity muscle use and ability to perform movements. Decreased muscle mass in the subcutaneous tissue is due to the accumulation of surrounding fatty tissue in the muscle, especially in elderly and obese individuals, resulting in a decrease in the size of type II muscle fibers [8]. Furthermore, in patients with serious medical conditions, decreased muscle mass on admission is associated with severity of illness, and decreased muscle quality is associated with an increased risk of infection and higher mortality in the intensive care unit [9]. These findings highlight the need for further investigation and countermeasures against muscle quality and quantity loss in children with physical disabilities due to cerebral palsy. Furthermore, evaluating how rehabilitation programs and exercise interventions can help improve muscle quality and function is crucial. A decrease in the percentage of muscle in the total subcutaneous tissue is characteristic of sarcopenia. Sarcopenia is caused by decreased activity and age-related reductions in testosterone and estrogen, leading to decreased muscle mass and accelerated fat accumulation [10]. The effects of adipocyte function and inflammation have also been reported, with hypertrophy of adipocytes in subcutaneous tissue being associated with inflammation and fibrosis progression [11]. For children with cerebral palsy, especially when verbal communication is challenging owing to the level of intellectual development, upper limb use becomes a primary mode of expression, such as using the upper limbs to guide and clap their hands. However, in these patients, muscle growth is impaired due to involuntary muscle contractions, resulting in decreased muscle size and tendon shortening [12]. Upper limb movements in children with cerebral palsy are often limited in movement patterns and speed, resulting in decreased accuracy [13]. Therefore, for participants in this study, the decreased percentage of muscle in the total subcutaneous tissue of the upper limb probably reflects inhibited muscle growth due to decreased upper limb activity and its effect on hormone secretion during the developmental stage.

Regarding the muscles of the lower limbs, this study revealed that the muscle thickness of the posterior aspect of the lower leg was significantly lower in children with physical disabilities and lesser mobility than in healthy children, suggesting that muscle strength in the lower limbs of healthy children is associated with developing functions related to many daily activities, especially those related to mobility involving the lower limbs. Muscle strength in the lower limbs is crucial in basic mobility skills such as walking and running, as well as more complex movements such as climbing stairs and jumping. Previous studies examining lower limb muscle strength and physical function in patients with cerebral palsy have demonstrated that ankle plantar flexor strength, in particular, is associated with functions such as walking distance and stair climbing, emphasizing the importance of lower limb muscle strength training in rehabilitating individuals with physical disability and cerebral palsy [14]. When implementing strength training, appropriate guidance should be provided to the patient, such as setting the appropriate amount of load and posture. However, communication is often challenging in children with physical disabilities owing to their level of intellectual development. Similarly, in this study, the intellectual development of children with physical disabilities ranged from those who can understand simple language to those who have difficulty understanding verbal communication, making it challenging to provide appropriate training instruction. For lower limb strength training in cerebral palsy, training programs targeting the antigravity muscles of the lower limbs in small groups (4–5 persons) have been proposed [15]. However, no specific programs for individuals who have difficulty communicating due to their level of intellectual development have been explicitly described in previous studies. A systematic review of the effectiveness of physical therapy for children with cerebral palsy revealed that while treatment interventions for the upper extremities were effective, strength training for the lower extremity muscles improved lower extremity muscle strength but not gait speed or stride length. There was also insufficient evidence that physical therapy interventions for the lower extremities improved movement ability [16]. This may be because providing a unified program for patients who have difficulty communicating is challenging due to their level of intellectual development. Therefore, physical therapy interventions are designed to increase muscle thickness in the lower limbs and improve movement performance by incorporating programs that encourage lower limb muscle activity in daily play and therapy activities. These programs are designed to increase the muscle thickness of the lower limbs and to improve movement ability. Among these, the evaluation of skeletal muscles using ultrasound imaging is expected to expand its use in clinical practice as a tool to measure outcome of physical therapy intervention for the lower limb because it can evaluate skeletal muscle thickness and structure noninvasively and realistically at a low cost and can track muscle changes before and after physical therapy intervention. Using ultrasound imaging for physical assessment will enable more accurate comprehension of skeletal muscle development in children with physical disabilities and support them in performing activities of daily living.

## Limitations

This study demonstrated the effectiveness of ultrasound imaging in the musculoskeletal assessment of children and adults with physical disabilities; however, several issues remain to be addressed. First, the limited sample size of the children and persons with disabilities included highlights the need for larger studies that include individuals with more diverse disabilities. Such studies will enable a more detailed analysis of the factors behind the decrease in muscle thickness and the development of rehabilitation programs for various types of disabilities. Another issue is the improvement in the accuracy of evaluation using ultrasound images and analysis techniques. Real-time evaluation is possible with ultrasound imaging; however, it is susceptible to noise caused by physical exercise, making it challenging to accurately evaluate skeletal muscle movement during actual physical activity. Previous studies evaluating ultrasonic imaging during physical exercise included an investigation of changes in the distance between the tibia and fibula when external rotation stress is applied to the foot by an external force [17]. Another study investigated the movement of the meniscus by temporally tracking the medial meniscus while walking [18] with an ultrasonic imaging device fixed to the knee with a band. However, these previous studies evaluated ultrasound images after fixing the limb position and direction of motion to some extent. In children with physical disabilities, specifying the direction of movement by instructions is often challenging owing to immobilization difficulties caused by involuntary movements or delayed intellectual development. Thus, dynamic evaluation using ultrasound images is challenging using the same method as in previous studies.

## Conclusions

This study highlights the importance of musculoskeletal assessment in rehabilitating children with physical disabilities and demonstrates the potential of using ultrasound imaging for assessment. Ultrasound imaging offers noninvasive, real-time assessment of musculoskeletal structures with minimal burden on the patient being evaluated. Decreased muscle thickness in children with physical disabilities may be due to multiple factors, such as decreased daily activity, decreased frequency of muscle use, or nutritional status. Further comprehensive studies are needed to clarify these factors. Future research holds promise for a more detailed understanding of functional impairment and the development of effective treatment and rehabilitation methods.

## Acknowledgments

We thank CosmoCare Energy, Inc. for its cooperation in the research of this study.

## Notes

### Competing Interest Statement

The authors have declared no competing interest.

